# Increase in sea-level and shelf area positively influenced the development of Cenozoic biodiversity hotspots by modulating turnover rate in marine benthos

**DOI:** 10.64898/2026.10.15.759295

**Authors:** Kella Venu Gopal, Devapriya Chattopadhyay

## Abstract

Marine biodiversity is unevenly distributed across the oceans, with exceptional concentrations of species richness confined to a few hotspots. Fossil and molecular evidence indicate that marine hotspots have shifted among ocean basins throughout the Cenozoic, but the mechanisms underlying their emergence and persistence have rarely been tested across multiple marine groups. Using standardized fossil occurrence data for molluscs, cnidarians, and foraminifera, we identified marine biodiversity hotspots through the past 66 million years. We then apply a multistep framework linking environmental drivers (sea surface temperature (SST), shelf area, and sea level) to clade-specific macroevolutionary processes (origination, extinction, and immigration) to evaluate the dynamics of these hotspots. Our results reveal contrasting evolutionary dynamics across marine biodiversity hotspots of different marine groups. Benthic groups: Mollusca and Cnidaria exhibit higher evolutionary turnover in hotspots than in non-hotspot regions, whereas planktic foraminifera maintain high diversity through reduced extinction and immigration rates. Increase in the sea level and shelf area positively contribute to the development of hotspot by increasing the origination rate in mollusca and cnidaria. Our study offers an integrative framework for understanding how marine biodiversity hotspots arise through a network of interacting processes: environmental drivers shape macroevolutionary dynamics, which in turn generate clade-specific diversity trajectories; together, these processes determine whether a particular region develops into a biodiversity hotspot.

**Significance Statement:** Biodiversity hotspots contain extraordinary concentrations of life, yet the evolutionary processes that generate them remain poorly understood in the oceans. Using a standardized Cenozoic fossil dataset, we reconstruct the emergence of marine biodiversity hotspots from major ocean basins. We find that hotspots exhibit elevated evolutionary turnover, but the underlying mechanisms vary among clades - ranging from rapid speciation to complex interactions among extinction, dispersal, and environmental change. These results show that marine hotspots arise through multiple evolutionary pathways rather than a single universal process, with important implications for conservation strategies.

## Introduction

Marine biodiversity is highly uneven across the world’s oceans, with exceptional concentrations of species richness confined to a few biodiversity hotspots (Myers et al., 2000; Myers, 2003). These regions play a disproportionate role in shaping global diversity patterns and have long been central to understanding large-scale ecological and evolutionary processes. Fossil and molecular evidence indicate that this spatial heterogeneity is not unique to the present day; marine biodiversity hotspots have appeared and disappeared across ocean basins through geologic time. In particular, Cenozoic records reveal a succession of major diversity centers across regions such as the Western Tethys, Arabian Sea, Indo-Australian Archipelago (IAA), and Caribbean - a pattern often described as the “Hopping Hotspot” model (Renema et al., 2008; Bellwood et al., 2012; Yasuhara et al., 2022). While these spatial shifts are well documented, the macroevolutionary mechanisms responsible for the emergence and persistence of marine biodiversity hotspots is not well resolved.

Several macroevolutionary hypotheses have been proposed to explain how hotspots form and maintain exceptionally high diversity. The Centre of Origin hypothesis proposes that hotspots function as cradles of diversification, generating new taxa (through increased rate of origination) that subsequently disperse to surrounding regions (Briggs, 2000; Gaither et al., 2013; Tornabene et al., 2015; Tian et al., 2024). In contrast, the Centre of Survival hypothesis suggests that hotspots act as long-term refugia with reduced extinction (Barber and Bellwood, 2005; Bellwood and Meyer, 2009). A third model, Centre of Accumulation, proposes that hotspots arise through the immigration and long-term accumulation of taxa originating elsewhere (Harzhauser et al., 2008; Leprieur et al., 2016; Huang et al., 2018; Miller et al., 2018; Yasuhara et al., 2022; Saulsbury and Baumiller, 2023). These hypotheses make contrasting predictions about the relative roles of origination, extinction, and immigration, providing a framework for distinguishing whether hotspots function primarily as cradles, refugia, or repositories of biodiversity. These macroevolutionary processes, in turn, is proposed to be modulated by environmental drivers such as sea-surface temperature (SST), tectonic reorganization, habitat heterogeneity, and the availability of shallow marine shelf area (Renema et al., 2008; Tittensor et al., 2010; Leprieur et al., 2016; Zaffos et al., 2017; Yasuhara et al., 2022; Tian et al., 2024). However, previous studies have largely focused on individual regions, taxonomic groups, or specific time intervals. The Indo-Australian Archipelago (IAA), one of the most prominent marine hotspots on Earth today, has been central to many of these debates. Previous studies on the IAA has proposed that its extraordinary diversity reflects combinations of high speciation, long-term persistence, and immigration from surrounding regions (Renema et al., 2008; Bellwood and Meyer, 2009; Cowman and Bellwood, 2013; Huang et al., 2018; Saulsbury and Baumiller, 2023; Tian et al., 2024). Yet it remains uncertain whether this region represents a general model for marine hotspot formation or whether different hotspots emerge through distinct evolutionary pathways. Resolving this question requires a broader comparative framework that evaluates hotspot dynamics across multiple clades, regions, and environmental contexts over deep time.

Here we identified major global marine biodiversity hotspots using a large fossil dataset of three major marine clades and standardizing it for sampling intensity through shareholder quorum subsampling. We evaluate the relative contributions of origination, extinction, and immigration to the persistence of hotspot. We further test if specific environmental variables (such as sea-surface temperature, shelf area, and sea-level change) functioned as drivers of hotspot formation by influencing specific macroevolutionary processes. Together, this framework allows us to test competing hypotheses for hotspot formation and assess the environmental and evolutionary dynamics underlying the emergence and persistence of marine biodiversity hotspots through deep time.

## Methods

### Data compilation and processing

We compiled a comprehensive database of Cenozoic marine fossil occurrences for three benthic groups: molluscs, benthic foraminifera, and cnidarians from the Paleobiology Database (PBDB; https://paleobiodb.org/; downloaded September 10, 2024), yielding a total of 235,661 occurrences. To improve geographic coverage in under sampled regions, we supplemented these data with 1,389 additional occurrences from published literature to improve coverage in understudied regions. Data for planktic foraminifera were obtained from the Triton dataset, comprising 1,805,688 occurrences (Fenton et al., 2021).

Only genus and species-level identifications were retained to ensure taxonomic reliability. We removed the records in which subgenera were entered in the genus field. Fossil occurrences were assigned to specific age-level time bins using the *bin_time*() function (method = “all”) in the *palaeoverse* R package. Based on these age assignments, we grouped the occurrences into Cenozoic sub-epochs for subsequent analyses. We also assigned the occurrences to four marine biogeographic regions: West Atlantic, Western Tethys, Northern Indian Ocean, and Indo-Australian Archipelago (IAA) (Fig. 1). We then calculated sub-epoch level fossil occurrences and genus richness for the four taxonomic groups in all the four biogeographic province.

**Figure 1:**
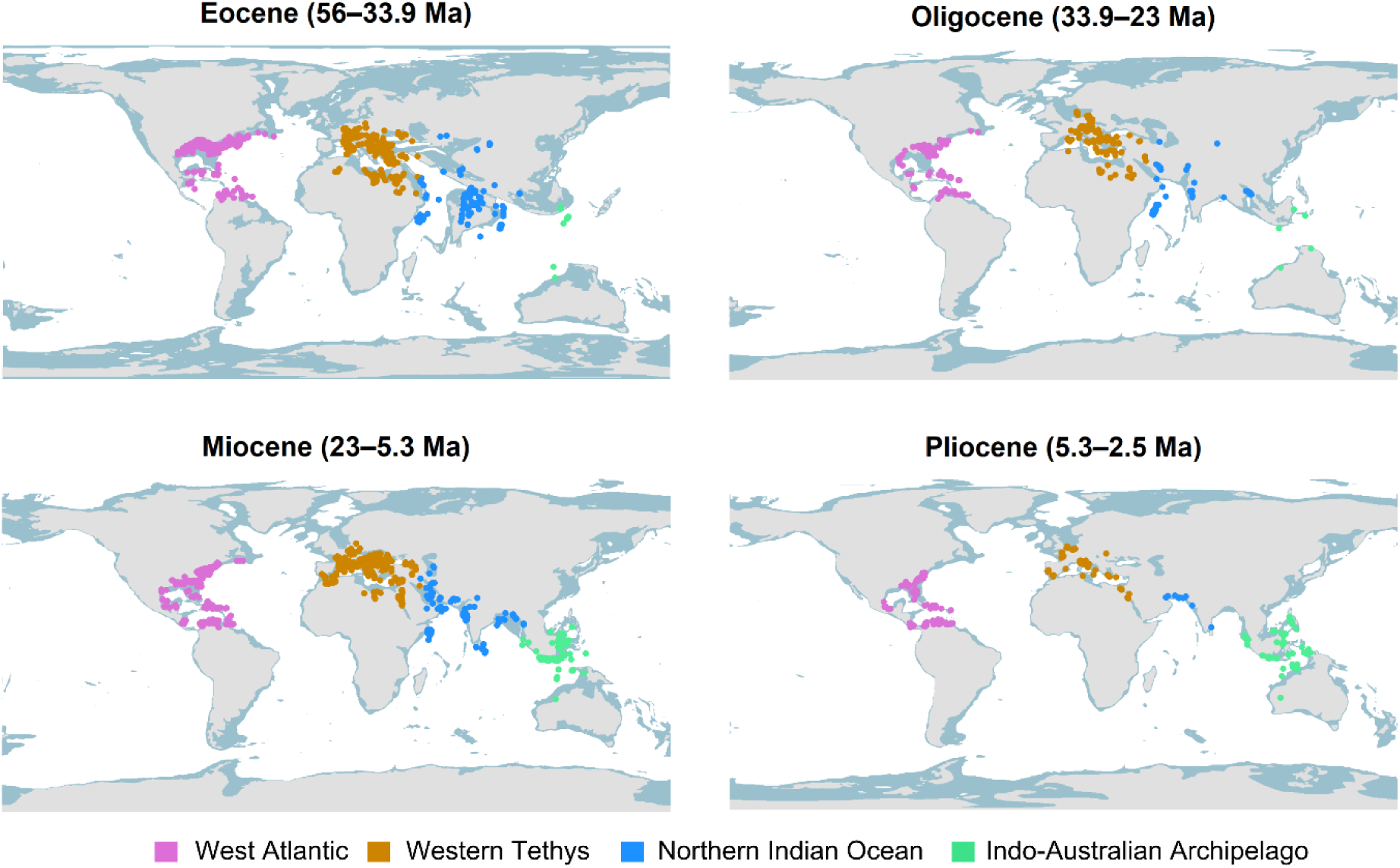
Spatial distribution of fossil occurrences for major marine groups across biogeographic provinces. Fossil occurrences of three major marine groups, Mollusca (open triangles), Cnidaria (open squares), and Benthic Foraminifera (open circles) across West Atlantic, Western Tethys, Northern Indian Ocean, and the Indo-Australian Archipelago. The data is sourced from PBDB.

### Sampling standardization and Hotspot identification

To assess the influence of sampling intensity on observed diversity patterns, we first examined the relationship between the number of fossil occurrences and observed genus richness. For each taxon–province combination, we calculated first differences in occurrence counts and genus richness between successive time bins and tested their correlation using Pearson correlation tests (Fig. S1). This approach evaluates whether temporal changes in observed richness are closely associated with changes in sampling intensity.

Genus richness was subsequently standardized using Shareholder Quorum Subsampling (Alroy, 2010) implemented in the *divDyn* R package (Kocsis et al., 2019). Sampling coverage was calculated for each taxon and province using the equation of Chao and Jost, (2012), and the median coverage across provinces was used to determine taxon-specific quorum levels. Quorum levels were 0.8 for Mollusca, 0.7 for benthic Foraminifera, 0.9 for planktic Foraminifera, and 0.8 for Cnidaria (Fig. S2). SQS diversity was estimated independently for the global dataset and each biogeographic province using 1,000 subsampling iterations, generating mean standardized genus richness and 95% confidence intervals (Section S1.2 of Supplementary). Following SQS standardization, we repeated the first-difference correlation analyses to evaluate whether residual sampling bias persisted in the standardized diversity curves (Fig. S3). This two-step procedure allowed us to assess the extent to which diversity patterns remained influenced by uneven fossil sampling after standardization. Regional and global standardized diversity curves were compared across the four biogeographic provinces for each taxon and sub-epochs. A biodiversity hotspot was defined when the mean standardized genus richness of a province was at least 50% of the corresponding mean global diversity for that time interval (Table 1).

**Table 1.** Temporal distribution of marine biodiversity hotspots across major Cenozoic biogeographic provinces. Hotspot occurrences are shown for four provinces—West Atlantic, Western Tethys, Northern Indian Ocean, and the Indo-Australian Archipelago— across sub-epochs. Hotspots are identified for four taxonomic groups: Mollusca (Mol), Cnidaria (Cni), benthic foraminifera (Bnf), and planktic foraminifera (Plf). Entries list the taxa for which a province qualifies as a hotspot within each interval. NH denotes that the province is not in a hotspot phase.

| Epoch_grad | West Atlantic | Western Tethys | Northern Indian Ocean | Indo-Australian Archipelago |
| --- | --- | --- | --- | --- |
| Paleocene | Bnf | Mol, Cni, Bnf, Plf | NH | NH |
| Early Eocene | NH | Cni, Bnf, Plf | NH | NH |
| Middle Eocene | NH | Mol, Cni, Plf | Plf | NH |
| Late Eocene | Bnf | Mol, Cni, Plf | Plf | NH |
| Early Oligocene | Cni | Bnf, Plf | Plf | NH |
| Late Oligocene | Bnf, Plf | Mol, Cni, Plf | Plf | NH |
| Early Miocene | Plf | Plf | Bnf, Plf | Cni |
| Middle Miocene | Plf | Bnf, Plf | Bnf, Plf | Cni |
| Late Miocene | Bnf, Plf | Bnf, Plf | Plf | Cni, Plf |
| Pliocene | Mol, Bnf, Plf | Bnf, Plf | Bnf, Plf | Cni, Plf |

### Diversity dynamics and standardization

We quantified three components of diversity dynamics (origination, extinction, and migration) for each taxon, province, and time bin. These components were measured using genus-level First Appearance Datum (FAD) and Last Appearance Datum (LAD) records, with genus ranges estimated from Phanerozoic fossil occurrence data. Global FAD and LAD ranges were used to identify origination and extinction events for each time bin. We defined the components as follows:

1. origination events: the number of genera whose FAD occurred within a given province and time bin;
2. extinction events: the number of genera whose LAD occurred within a given province and time bin;
3. migration events: the first occurrence of a genus in a province other than its inferred province of origin.

The province of origin was assigned based on the earliest occurrence of a genus across the four study provinces. For migration, only genera that had originated previously in another province were considered, and only their first occurrence in the new province was counted as a migration event. This approach minimized the potential influence of repeated occurrences of the same genus following its initial extra-provincial appearance. We compared the relative contributions of origination, extinction, and migration between hotspot and non-hotspot provinces. Because differences in fossil sampling intensity among provinces could influence estimates of these macroevolutionary processes, we further standardized the process-based analyses using a province-level resampling procedure. Using the taxon-specific coverage-based quorum used for SQS diversity standardization, we randomly sampled the occurrence data with replacement from each province (Section S2.1 of Supplementary). The resampled records from all four provinces were then used to identify origination, extinction, and migration events. This procedure was repeated 1,000 times for each taxon. Mean and standard deviation of the number of genera assigned to each process were calculated for every province–time-bin combination. The mean process counts were subsequently normalized by the total number of fossil occurrences to account for differential sampling intensity in each province–time combination (Section S2.1 of Supplementary), yielding occurrence-standardized estimates of origination, extinction, and migration (Table S3).

### Environmental predictors

Region-specific SST reconstructions were not available for all provinces; SST values were assigned according to broad climatic zones. Temperate SST values were assigned to the Western Tethys, whereas tropical SST values were assigned to the West Atlantic, Northern Indian Ocean, and Indo-Australian Archipelago (IAA). The West Atlantic was classified as a tropical region because most fossil occurrences from this province are concentrated in the Caribbean, a region recognized as a tropical hotspot (Yasuhara et al., 2022).

Global sea-level fluctuations were obtained from the smoothed sea-level curve of Miller et al., (2020), derived from Pacific benthic foraminiferal δ¹⁸O and Mg/Ca records. Sea-level values were assigned to each Cenozoic sub-epoch by calculating the mean smoothed sea level within a ±1 Myr window around the midpoint age of each time interval. The resulting shelf-area, SST, and MSL estimates for province-time bin combinations were used as environmental predictors in subsequent analyses.

### Statistical analysis

We evaluated the hierarchical relationships linking environmental variables, macroevolutionary processes, and hotspot occurrence, we implemented a Bayesian structural equation model (SEM) using the *brms* package in R (Bürkner, 2017). Three Gaussian sub-models were specified to quantify the effects of SST, shelf area, and sea level on origination, extinction, and migration rates. Hotspot occurrence (presence/absence) was modelled using a Bernoulli distribution with origination, extinction, and migration as predictors. We included taxon as a varying-effects grouping factor with random intercepts and slopes, allowing both environmental effects on macroevolutionary processes and the influence of these processes on hotspot probability to vary among clades. Residual correlations among the process models were not estimated to preserve directed causal pathways. Models were fitted using four Markov chain Monte Carlo method (10,000 iterations each, 200 warm-up) with weakly informative priors and stringent sampling controls (*adapt_delta* = 0.995; *max_treedepth* = 18). Convergence was confirmed using R^ values (∼1.0) and adequate effective sample sizes. We calculated path coefficients for each taxon as the sum of global fixed effects and taxon-specific deviations (Table S4). We estimated Indirect (mediated) environmental effects on hotspot occurrence as the product of (i) the environment– process slope and (ii) the process–hotspot slope on the logit scale (Table S5).

## Results

### Cenozoic distribution of marine biodiversity hotspots

Cenozoic fossil occurrences of molluscs, cnidarians, and foraminifera cluster into four major marine provinces: West Atlantic, Western Tethys, Northern Indian Ocean, and Indo-Australian Archipelago (IAA) (Fig. 1). Raw genus richness varies substantially among provinces, taxa, and time intervals (Fig. 2a-d). Across most datasets, raw richness is strongly correlated with the number of fossil occurrences, indicating that sampling intensity influences observed diversity patterns (Fig. S1). However, these significant correlations disappear when raw occurrence counts were compared against SQS standardized diversity curves, except for molluscs and cnidarians from the Northern Indian Ocean (Fig. S3). Therefore, the standardized regional and global diversity curves were generated after excluding them (Fig. 2e-h).

**Figure 2.**
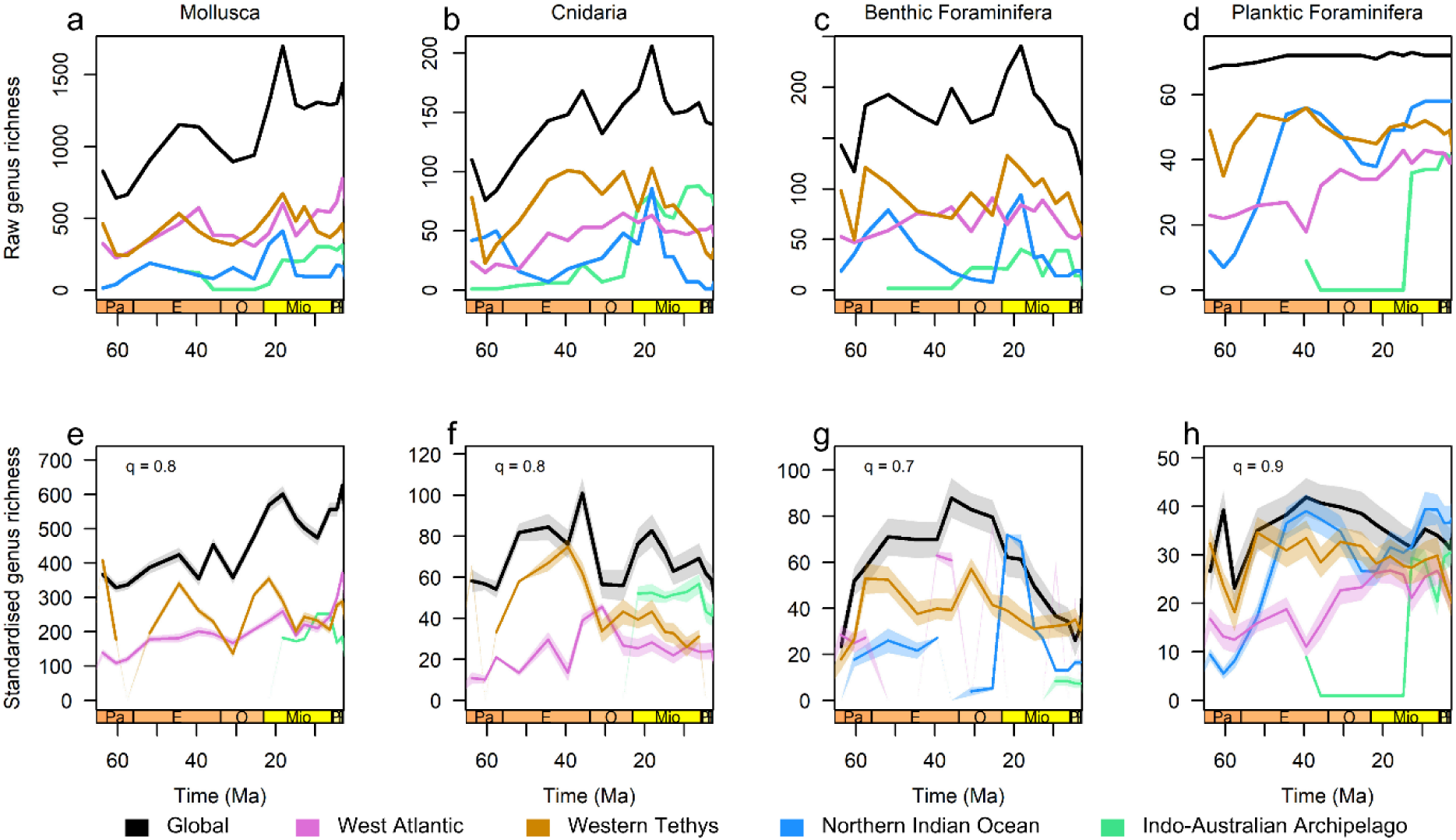
Global and regional diversity trends across the Cenozoic for four major marine groups - Mollusca, Cnidaria, Benthic Foraminifera, and Planktic Foraminifera, across four biogeographic provinces: West Atlantic, Western Tethys, Northern Indian Ocean, and Indo-Australian Archipelago. (a-d): Raw genus richness curves. (e-h): Standardised diversity curves generated using SQS method, with applied quorum levels (Mollusca = 0.8, Cnidaria = 0.8, Benthic Foraminifera = 0.7, and Planktic Foraminifera = 0.9).

Standardized diversity trajectories of provinces do not consistently track global diversity trends (Fig. 2e-h). Provinces gained and lost their hotspot status with varying contribution from different taxonomic group through time (Table 1, Fig. S7). Western Tethys represented the most consistently diverse multi-clade hotspot during the Paleocene–Early Miocene, whereas later Cenozoic hotspot occurrence became increasingly dominated by foraminiferal groups across the Western Tethys, Northern Indian Ocean, and West Atlantic, and by cnidarians and planktic foraminifera in the Indo-Australian Archipelago.

### Macroevolutionary dynamics of hotspot regions

The difference in macroevolutionary processes between hotspot and non-hotspot provinces varies across taxonomic groups (Fig. 3). Molluscan and cnidarian hotspots exhibit increased rates of both origination and extinction, resulting in higher overall turnover (Fig. 3a-b). Benthic foraminifera do not show any significant difference (Fig. 3c). In contrast, planktonic foraminiferal hotspots are associated with significantly lower extinction and immigration rates than non-hotspot regions (Fig. 3d).

**Figure 3.**
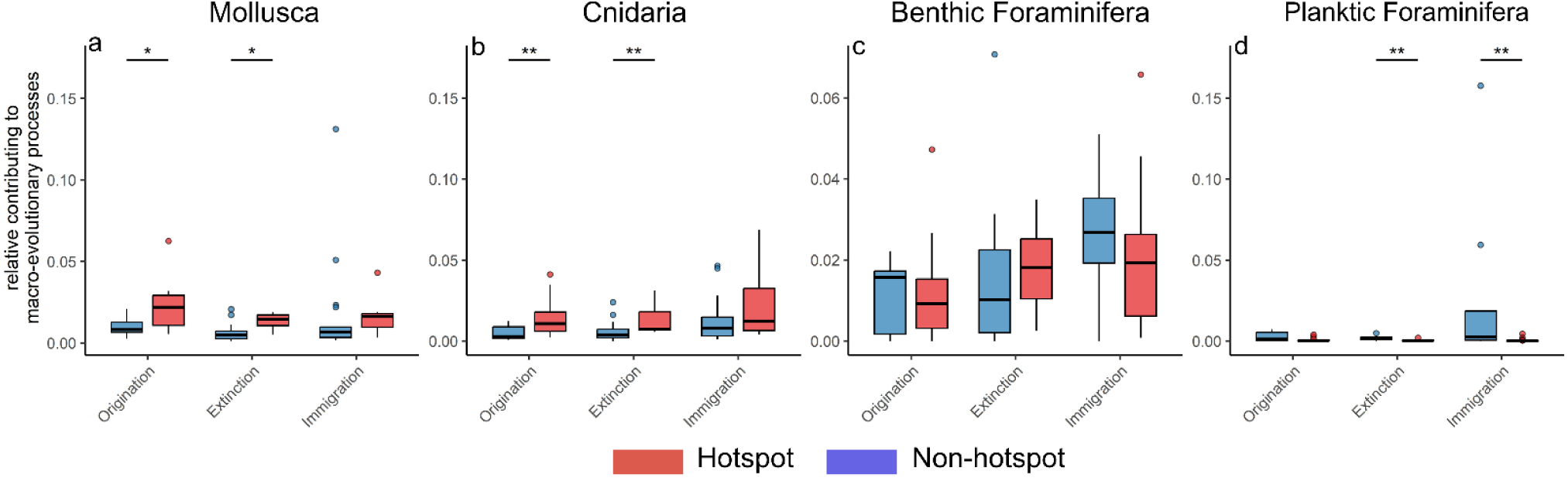
Comparison of the number of genera contributing to macroevolutionary processes in hotspot versus non-hotspot provinces. Panels show results for Mollusca (a), Cnidaria (b), benthic foraminifera (c), and planktic foraminifera (d). Boxplots illustrate the distributions of origination, extinction, and immigration in hotspot (red) and non-hotspot (blue) provinces. Horizontal bars with asterisks indicate statistically significant differences between groups (* p < 0.05, ** p < 0.01, *** p < 0.001).

### Environmental controls on macroevolutionary processes

The causal networks reveal strong clade-specific connection between the environmental and macroevolutionary controls of hotspot dynamics (Fig. 4). In molluscs and cnidarians, shelf area and mean sea level exhibit significant positive effects on origination, which in turn shows a strong positive association with hotspot occurrence (Fig. 4). In benthic and planktic foraminifera, none of the environmental variables show significant association with any of the macroevolutionary processes (Fig. 4). SST shows predominantly non-significant relationships with macroevolutionary processes in all clades.

**Figure 4.**
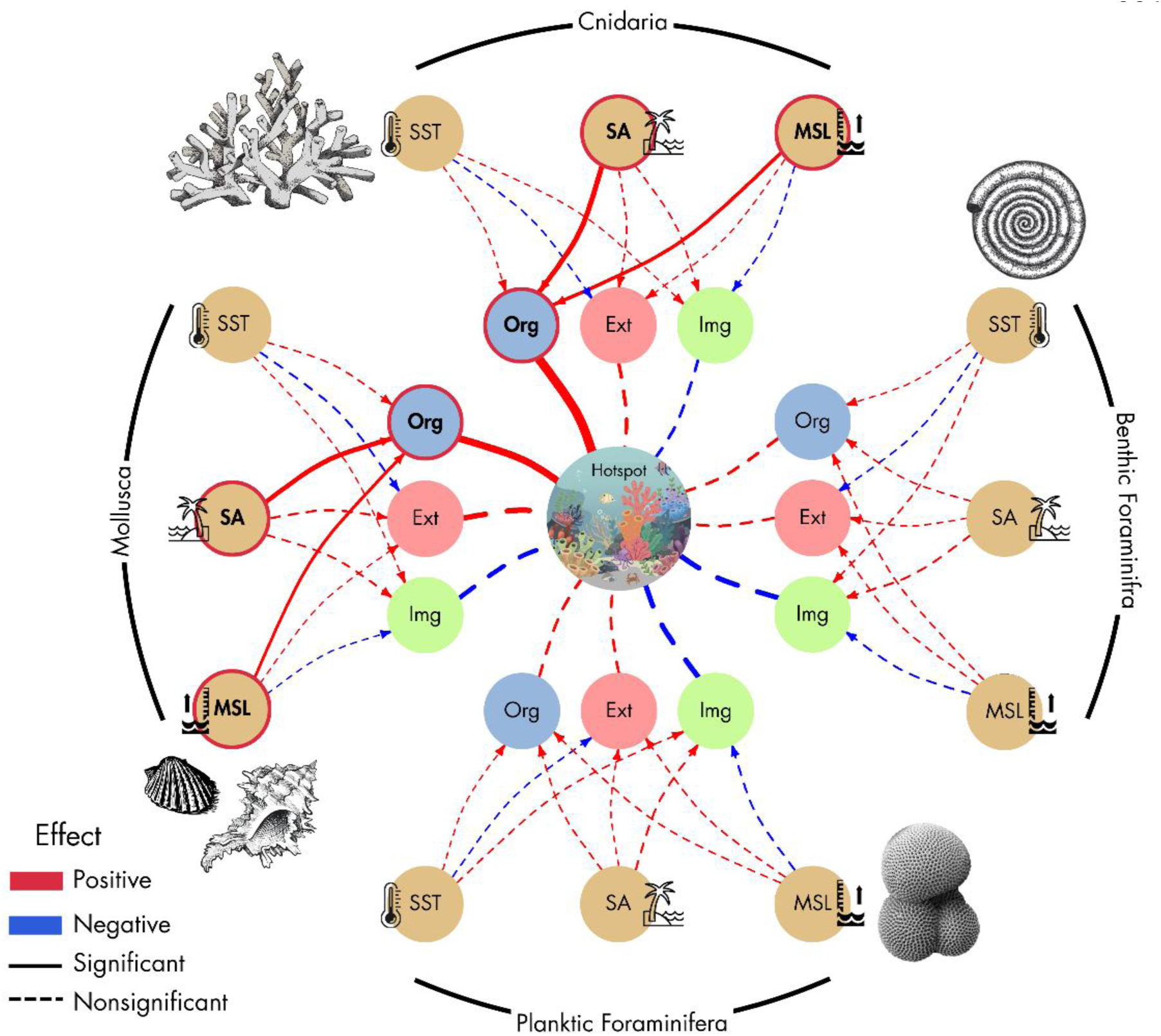
Causal network diagrams illustrating environmental–process–hotspot pathways for major marine taxa. Networks depict relationships among environmental variables (SST, sea surface temperature; SA, shelf area; MSL, mean sea level), macroevolutionary processes (Org, origination; Ext, extinction; Img, immigration), and hotspot dynamics for Mollusca, Cnidaria, benthic foraminifera, and planktic foraminifera. Arrows indicate causal links, with edge thickness proportional to effect size and colour denoting direction (red, positive; blue, negative). Solid lines represent statistically significant effects, whereas dashed lines indicate non-significant links.

## Discussion

### Evolutionary dynamics within marine biodiversity hotspots through time

Molluscan and cnidarian hotspots exhibit a significantly higher origination and extinction rates (Fig. 3a-b), indicating elevated evolutionary turnover rather than simple long-term taxonomic accumulation. Such elevated evolutionary turnover parallels patterns documented in terrestrial biodiversity hotspots, where rapid lineage replacement has been proposed as an important mechanism underlying the maintenance of exceptional diversity (Ge et al., 2022; Sonne and Rahbek, 2024). Several studies have suggested that the IAA acts as a cradle of speciation, generating new taxa that subsequently disperse to surrounding regions (Briggs, 2000; Gaither et al., 2013; Tornabene et al., 2015; Tian et al., 2024). Our results, although consistent with higher origination in molluscan and cnidarian hotspots, highlights the role of elevated extinction. Similar patterns of relatively high coral extinction have been reported from the Coral Triangle (Huang et al. 2018), suggesting that origination and extinction may operate simultaneously within regions of exceptionally high diversity. In contrast to the premise of Centre of Accumulation model, immigration was not found to play an important role in the development of molluscan and cnidarian biodiversity hotspots (Fig. 3a-b). This observation also contrasts previous studies proposing that immigration and regional species accumulation played an important role in the development of the modern IAA biodiversity hotspot across multiple benthic taxa (Harzhauser et al., 2008; Mihaljević et al., 2017; Huang et al., 2018; Yasuhara et al., 2020a; Saulsbury and Baumiller, 2023).

Planktic foraminiferal hotspots in our study exhibit significantly lower immigration and extinction than non-hotspot regions, whereas origination does not differ significantly (Fig. 3d). The reduced immigration is consistent with previous work suggesting that diversification in planktic foraminifera is strongly associated with ecological partitioning along environmental gradients rather than dispersal alone (Seears et al., 2012). Closely related taxa frequently differentiate along environmental gradients in sea-surface temperature, mixed-layer depth, thermocline structure, salinity, and seasonality, allowing multiple species to coexist within the same broad geographic region through niche differentiation (Fenton et al., 2016; Antell et al., 2021; Mathes et al., 2024). In such systems, hotspot regions may already contain a dense assemblage of specialised taxa occupying available ecological niches. Under these conditions, environmental filtering and incumbency effects may limit the successful establishment of immigrant species, resulting in relatively low immigration rates. This suggests that, for pelagic groups, a hotspot is a Centre of Survival where the diversity is maintained primarily through in situ ecological diversification and low extinction, rather than through continuous influx from surrounding regions. The observed consistency of macroevolutionary dynamics among benthic groups and its deviation from the planktic groups are also expected. Marine benthic macroevolutionary dynamics are decoupled from planktonic group dynamics, reflecting distinct evolutionary drivers between benthic and planktonic marine realms (Clarke and Crame, 2010; Erwin, 2024). Benthic groups show long-term steady-state diversity despite high turnover (Jablonski and Sepkoski, 1996), in contrast to planktonic groups that display faster turnover tightly coupled to surface-ocean productivity, nutrient stress, and climate fluctuations (Alvarez et al., 2019; Tingle et al., 2025).

### Environmental mediated pathways to marine biodiversity hotspots

Our results indicate that environmental triggers do not directly translate into marine biodiversity hotspots, but instead influence hotspot formation by modulating specific macroevolutionary pathways (Fig. 4). In molluscs and cnidarians, increased shelf area and mean sea level were positively associated with origination, which in turn was positively associated with hotspot occurrence (Fig. 4, Table S5). This pathway suggests that expansion of shallow-marine habitat may have increased the availability of ecological space and opportunities for lineage establishment, thereby contributing to the development of regional diversity concentrations. Importantly, extinction was not significantly associated with the environmental variables examined here, indicating that the environmental triggers primarily impacted origination rather than influencing lineage loss. Similarly, immigration did not provide a significant environmental pathway to hotspot occurrence in any of the groups.

The association between expanded shallow-marine habitat and origination is consistent with the long-standing recognition that periods of high sea level and extensive shelf increase the availability and heterogeneity of marine habitats. As rising seas inundate continental shelves, accommodation space for shallow marine habitats (reef, seagrass, mangrove) expands and contribute heavily in developing biodiversity hotspots (Graham et al., 2003; Gordillo et al., 2010; Aguirre et al., 2011; O’Dea et al., 2026). Greater habitat area and environmental heterogeneity can provide additional opportunities for ecological differentiation and lineage establishment, potentially contributing to pulses of origination and regional diversity (Smith and Benson, 2013; Leprieur et al., 2016; Zaffos et al., 2017; Harzhauser et al., 2024; Venu Gopal et al., 2025). A reduction in shelf area, unlike origination, does not consistently drive extinction in marine benthos (Clarke et al., 2004). Benthic extinction is more often driven by changes in ocean circulation and habitat fragmentation (Thatje et al., 2005; Bottjer et al., 2008; Clarke and Crame, 2010) – parameters that we did not consider in our environmental variables. This explains why extinction, despite differing significantly between hotspot and non-hotspot comparisons (Fig. 3), does not emerge as an important macroevolutionary mechanism linking environmental parameters to hotspot development (Fig. 4).

Across the four clades, SST shows predominantly non-significant relationships with macroevolutionary processes (Fig. 4). These findings contrast with several studies that identify SST as major correlate of present day marine biodiversity hotspots (Tittensor et al., 2010; Fenton et al., 2016; Saeedi et al., 2017; Dalongeville et al., 2022; Förderer et al., 2023). This discrepancy likely reflects the Cenozoic temperature variability which remained largely within the physiological tolerances of many marine groups, thereby reducing its impact on origination and extinction dynamics. Moreover, the SST variable used here represents broad temporal and spatial environmental conditions. Such coarse resolution may not capture the ecological variables that directly regulate the distribution or evolutionary dynamics of individual clades.

The absence of significant environmental pathways in foraminifera contrasts with the molluscan and cnidarian patterns. Even though, planktic foraminifera showed reduced extinction and immigration for hotspot in contrast to non-hotspot regions (Fig. 3), none of the foraminiferal groups demonstrated significant association with the environmental triggers (Fig. 4). The observed patterns in Mollusca and Cnidaria are unlikely to be artefacts of geological heterogeneity (Section S1.3 of Supplementary). Both groups show relatively homogeneous lithological composition among provinces (Table S1, Fig. S5) and consistently low dissimilarity in depositional environment (Table S2, Fig. S6). This absence suggests that hotspot dynamics in foraminifera may be governed by factors outside our present models. Benthic foraminifera showed the largest differences among provinces in both lithological composition (Table S1, Fig. S5) and depositional environment (Table S2, Fig. S6). Modern benthic foraminiferal diversity, for example, is shaped mainly by depth, dissolved oxygen, facies type, and only weakly by the shelf area (Smith et al., 2001; Smith and Gallagher, 2003; Goeting et al., 2023). Macroevolutionary dynamics of benthic foraminifera is also decoupled from the largely climate-driven Cenozoic diversity trends seen in planktonic groups. Their diversity instead tracks local water-mass, oxygenation, and productivity changes rather than the global climate signal that shape the diversity of planktonic foraminifera (Yasuhara et al., 2020). Alternatively, the comparatively high dispersal potential of foraminifera, particularly in planktonic taxa, may have weakened the link between provincial environmental conditions and the origination or immigration measures used here.

## Conclusion

Our study offers an integrative approach to understand how marine biodiversity hotspots arise through a network of interacting processes: environmental drivers shape macroevolutionary dynamics, these dynamics generate clade-specific diversity trajectories, and together they determine the likelihood that a particular region develops into a biodiversity hotspot. Benthic groups, mollusca and cnidaria, maintain diversity in hotspots through high rate of turnover, in contrast to planktic foraminifera that maintains diversity through low extinction. Increase in the sea level and shelf area positively contribute to the development of hotspot by increasing the origination rate in mollusca and cnidaria.

### Reproducibility

All data and analysis scripts are publicly available through a GitHub repository (https://github.com/VenuGopalK08/Cenozoic-hotspot-dynamics.git). The repository contains separate directories for each analytical component, including associated datasets, code, and documentation. Analyses were conducted using R (v. 4.2.2) on Windows 11 systems.

## Data Availability Statement

All relevant data supporting used for our analyses are available on the associated GitHub repository (https://github.com/VenuGopalK08/Cenozoic-hotspot-dynamics.git).

## Supporting information

Supplementary Analysis, Figures and Tables

## Acknowledgements

We thank all contributors to the Paleobiology Database, without whom this work would not be possible.

## Authors contributions

KVG and DC conceived and designed the project and led the writing of the manuscript. KVG curated the data, conducted the analyses, prepared figures.

