## Supplementary Analysis, Figures and Tables for "Increase in sea-level and shelf area positively influenced the development of Cenozoic biodiversity hotspots by modulating turnover rate in marine benthos"

Kella Venu Gopal[1*] and Devapriya Chattopadhyay[1]
[1]IISER Pune, Dr. Homi Bhabha Road, Pashan, Pune – 411008

* Corresponding author

This pdf contains:
Section: S1 to S4
Fig. S1 to S7
Table S1 to S5

**INDEX**

S1. Standardization of occurrence data

*S1.1. Heterogeneous sampling across taxa and provinces*

*S1.2. Taxon-specific standardization using Shareholder Quorum Subsampling (SQS)*

*S1.3. Assessing the geological and preservation biases among different provinces*

S2. Standardization of macroevolutionary processes

*S2.1. Resampling the occurrence data using Coverage-quorum (q-value) resampling*

*S2.2. Resampling the occurrence data using minimum occurrence rarefaction resampling*

*S2.3. Dominant macroevolutionary processes operating in hotspots*

S3. Evaluating environmental variation across hotspot and non-hotspot region

*S3.1. Sea surface temperature*

*S3.2. Change in regional shelf area*

*S3.3. Global mean sea level*

S4. Environmental and macroevolutionary drivers of Cenozoic Hotspot Dynamics

*S4.1. Direct effects*

*S4.2. Indirect effects*

### S1. Standardization of occurrence data

#### *S1.1. Heterogeneous sampling across taxa and provinces*

We wanted to quantify which provinces and taxonomic groups are affected by sampling biases. Therefore, we performed first-difference correlations between fossil occurrences (proxy for sampling intensity) and genus richness to understand this effect (Fig. S1). Mollusca, Cnidaria, and benthic foraminifera exhibit strong positive correlations between changes in

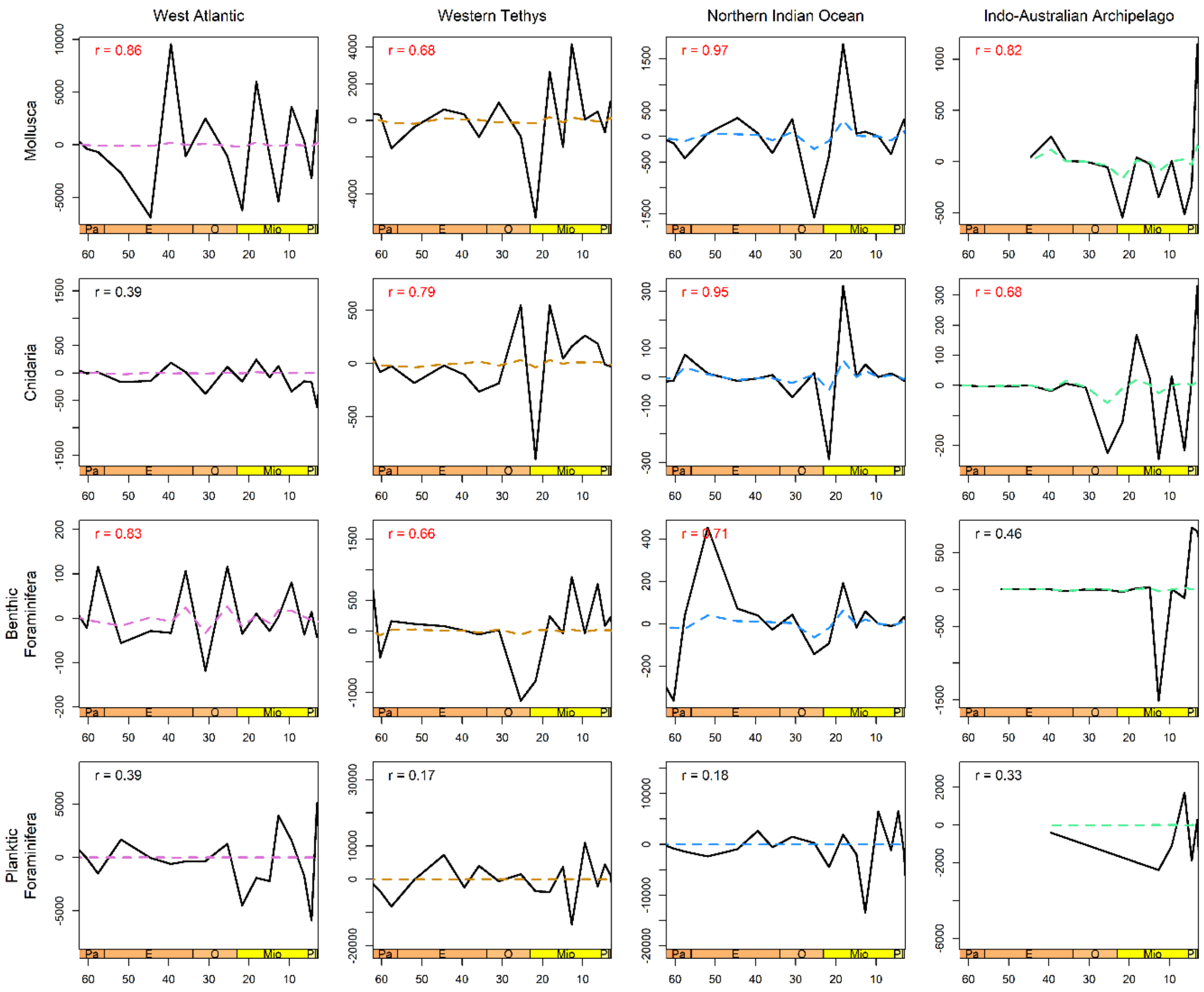


**Figure S1:** First-difference correlation analyses between raw genus richness and fossil occurrence counts across the Cenozoic for four major taxonomic groups (Mollusca, Benthic Foraminifera, Planktic Foraminifera, and Cnidaria) across four biogeographic provinces: West Atlantic, Western Tethys, Northern Indian Ocean, and Indo-Australian Archipelago. The solid black lines represent first differences in fossil occurrence, while colored dashed lines represent first differences counts sampled-in-bin genus richness. Pearson correlation coefficients (r) are shown in each panel, with significant correlations ($p < 0.05$) in red.

richness and occurrences across most provinces, suggesting that sampling biases partly contribute to their observed diversity fluctuations. In contrast, planktic foraminifera shows limited influence of sampling on their diversity patterns.

*S1.2. Taxon-specific standardization using Shareholder Quorum Subsampling (SQS)*

We applied Shareholder Quorum Subsampling (SQS) (Alroy, 2010) to generate regional and global standardized diversity curves. We used equation provided by Chao and Jost, (2012) for calculating the respective median coverage (q) values over entire Cenozoic Era for each taxonomic group: Mollusca = 0.8, Cnidaria = 0.8, Benthic Foraminifera = 0.7, and Planktic Foraminifera = 0.9 (Fig. S2).

Chao and Jost (2012) provide the following equation for coverage (q):

$$q = 1 - (f1 / n) * [ ((n - 1) * f1) / ((n - 1) * f1 + 2 * f2)]$$

- q = Sample coverage (completeness of sampling)
- n = Total number of individuals (or occurrences) in the sample
- $f_1$ = Number of singleton taxa (taxa represented by only one occurrence)
- $f_2$ = Number of doubleton taxa (taxa represented by exactly two occurrences)

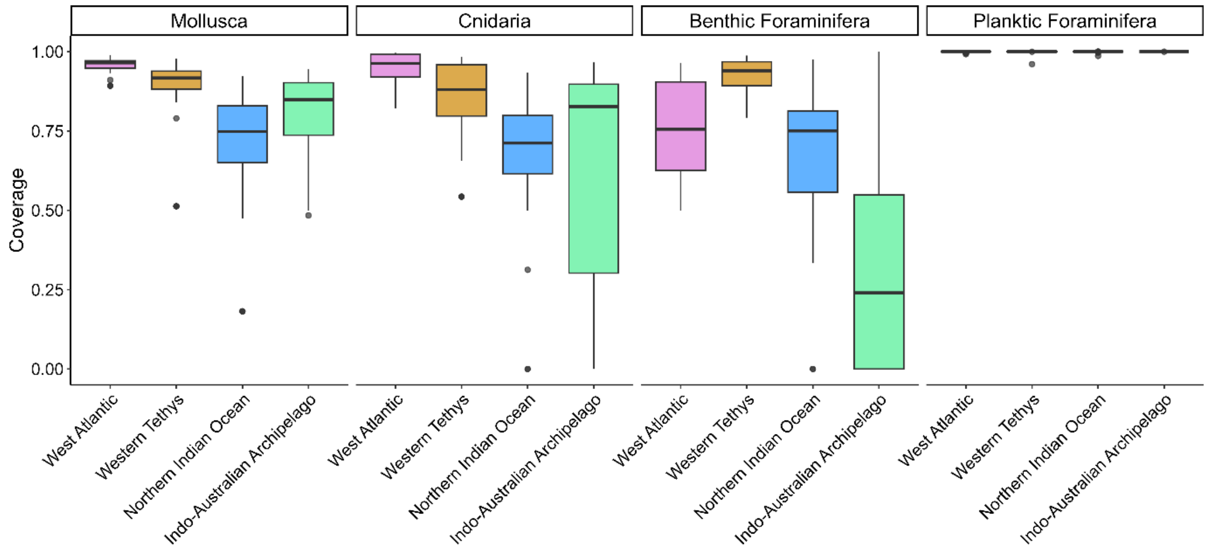


**Figure S2:** Median coverage (q) values for marine taxa across the ocean basins during Cenozoic.

We applied SQS standardization to raw genus richness and re-ran first-difference correlation tests between standardized genus diversity and occurrence counts to assess whether residual sampling bias persisted (Fig. S3). SQS substantially reduced the influence of sampling intensity, as the correlation between occurrence counts and standardized diversity largely disappeared after subsampling (Fig. S3). However, Mollusca and Cnidaria in the Northern Indian Ocean continued to show a strong dependence of diversity on sampling intensity (Fig. S3), limiting the reliability of SQS standardization for these datasets. Therefore, we removed the Mollusca and Cnidaria data from the Northern Indian Ocean from our hotspot analysis (Fig. 2e-f). In addition, several time bins for certain taxa could not produce standardized diversity curves because the required coverage thresholds were not met (Fig. 2e-f). These temporal bins and provinces were therefore not considered for hotspot analysis.

SQS standardization can nevertheless be sensitive to highly uneven assemblages, particularly under low sampling intensity (Alroy, 2010; Close et al., 2018). This is especially relevant for hotspot identification, as hotspots may contain a high proportion of rare taxa. Therefore, standardization to equal coverage across provinces provides a conservative estimate of diversity by reducing the influence of differences in sampling intensity among regions and time bins. Consequently, the hotspot patterns identified after SQS standardization are less likely to be driven solely by elevated sampling intensity. However, SQS cannot completely remove biases associated with the heterogeneous geological record, as differences in lithology, depositional environment, rock record, preservation, and exposure can still influence observed diversity.

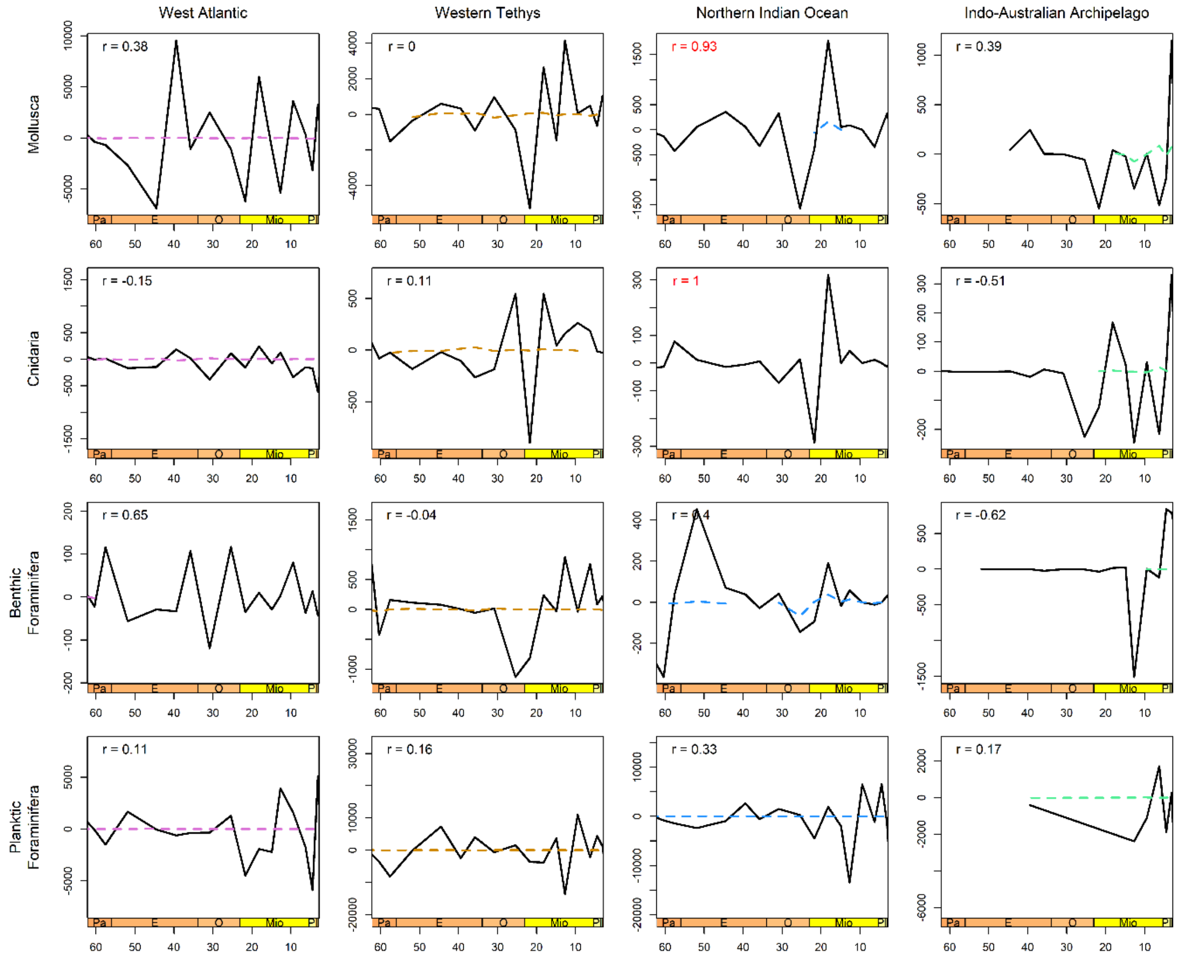


**Figure S3:** First-difference correlation analyses between SQS standardized genus richness (quorum levels: Mollusca = 0.8, Cnidaria = 0.8, Benthic Foraminifera = 0.7, and Planktic Foraminifera = 0.9) vs fossil occurrence counts (Age level) across provinces for four major shallow marine groups. The solid black lines represent first differences in fossil occurrence, while coloured dashed lines represent first differences of standardized genus richness. Pearson correlation coefficients (r) are shown in each panel, with significant correlations ($p < 0.05$) in red.

*S1.3. Assessing the geological and preservation biases among different provinces*

*Differential rock record among different provinces*

We assessed the biases of the geological record independently of occurrence abundance in each province by studying the number of unique stratigraphic formations represented in each taxon-province-time interval combination (Fig. S4). Formation names reported from PBDB and were treated as discrete stratigraphic units, and the number of unique formations therefore provides a measure of the breadth of stratigraphic units contributing fossil occurrences to each taxon, province, and time interval.

Formation counts provide an indication of the breadth of documented stratigraphic units contributing fossil occurrences, but do not directly measure sedimentary rock volume, preservation quality, or sampling intensity. Accordingly, we interpreted formation counts as a proxy for geological representation rather than as an unbiased measure of sampling completeness. The number of occurrences assigned to different lithological categories was highest in the West Atlantic and Western Tethys across most taxonomic groups (Fig. S4). Both Northern Indian Ocean and IAA showed considerable lower formations counts for all taxa across most Cenozoic intervals.

The substantially greater number of geological formations in the Western Tethys and West Atlantic indicates greater geological sampling opportunity in these provinces, whereas the sparse record of the IAA, particularly during the Paleogene, indicates substantially lower sampling opportunity. Consequently, spatial and temporal differences in observed diversity and evolutionary dynamics should be interpreted in the context of heterogeneous geological representation.

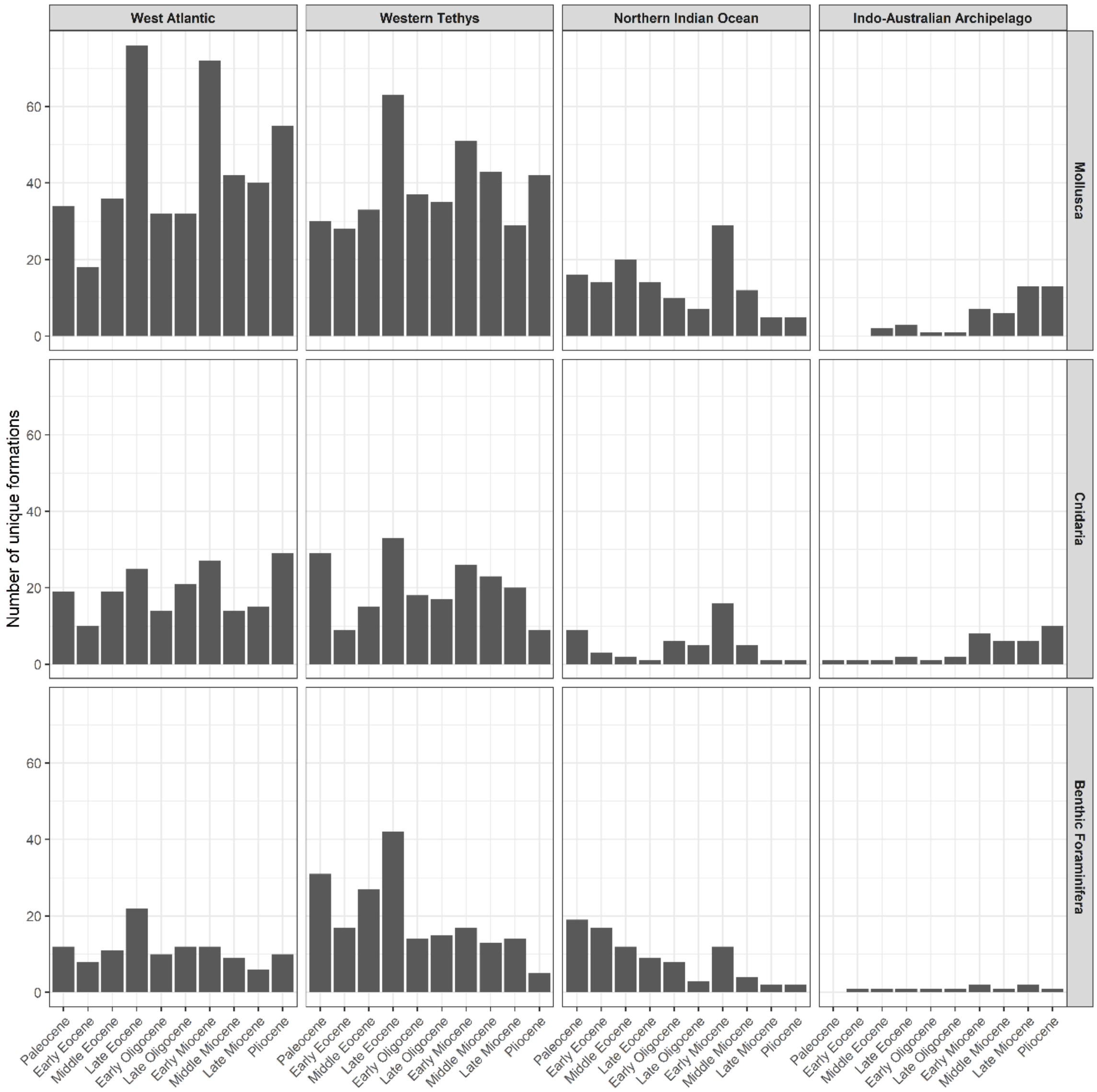


**Figure S4.** Temporal distribution of unique stratigraphic formations across four biogeographic provinces (West Atlantic, Western Tethys, Northern Indian Ocean, and Indo-Australian Archipelago) for Mollusca, Cnidaria, and benthic Foraminifera throughout the Cenozoic.

*Lithological differences between provinces*

The PBDB and literature based lithology were broadly grouped into six major groups. *Carbonate* lithologies included limestone, carbonate, chalk, dolomite, lime mudstone, grainstone, packstone, wackestone, rudstone, floatstone, framestone, bafflestone, bindstone, and reef rocks. *Siliciclastic* lithologies included sandstone, siltstone, mudstone, claystone, shale, conglomerate, gravel, breccia, and quartzite. *Mixed carbonate-siliciclastic* lithologies included marl and mixed carbonate-siliciclastic deposits. *Biogenic/chemical* deposits include chert, diatomite, coal, peat, ironstone, tar and phosphorite. *Volcaniclastic* includes ash, tuff, and volcaniclastic deposits.

Lithological composition varied among provinces and taxonomic groups across the Cenozoic (Fig. S5). The most distinct lithological combinations among provinces were identified using Bray-Curtis similarity threshold of 0.5 (0 indicate greater similarity and values closer to 1 indicate dissimilarity) (Table S1). Mollusca showed relatively similar lithological composition among all four provinces, with Bray-Curtis dissimilarity values ranging from 0.11 to 0.22, and none of the province pairs crossed the 0.5 dissimilarity threshold. Cnidaria showed greater differences in lithological composition, with the highest dissimilarity observed between the West Atlantic and Indo-Australian Archipelago (0.51; similarity = 0.49). Benthic Foraminifera showed the greatest overall lithological differences among provinces, with dissimilarity values ranging from 0.21 to 0.67. The West Atlantic and Indo-Australian Archipelago showed the highest dissimilarity (0.67; similarity = 0.33), representing the most distinct lithological combination in the dataset.

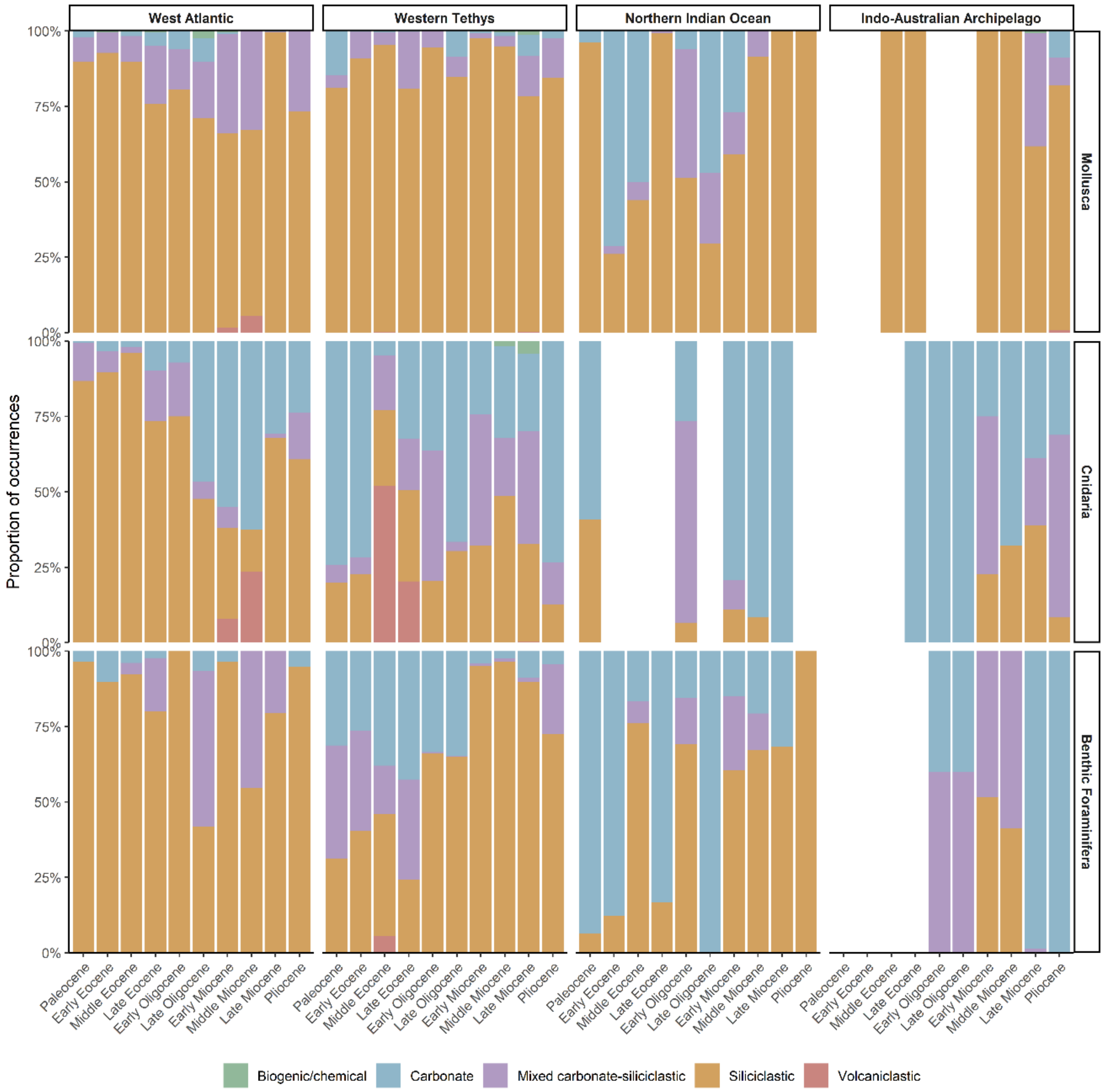


**Figure S5.** Lithological composition of fossil occurrences across Cenozoic marine biodiversity provinces. Proportions of occurrences assigned to major lithological categories are shown for Mollusca, Cnidaria, and benthic Foraminifera across the West Atlantic, Western Tethys, Northern Indian Ocean, and Indo-Australian Archipelago through successive Cenozoic epochs. Occurrences with unreported lithologies are excluded from the analysis.

**Table S1.** Bray-Curtis dissimilarity matrices showing differences in the composition of lithology among the four biogeographic provinces for Mollusca, Cnidaria, and Benthic Foraminifera across the Cenozoic. Occurrences with unreported lithology were excluded from the analysis. *Cnidaria and Mollusca were removed from Northern Indian Ocean.

| **Taxon** | Province | West Atlantic | Western Tethys | Northern Indian Ocean | Indo-Australian Archipelago |
|---|---|---|---|---|---|
| Mollusca | West Atlantic | 0.00 | | | |
| | Western Tethys | 0.11 | 0.00 | | |
| | Northern Indian Ocean | NA* | NA* | 0.00 | |
| | Indo-Australian Archipelago | 0.22 | 0.22 | NA* | 0.00 |
| Cnidaria | West Atlantic | 0.00 | | | |
| | Western Tethys | 0.38 | 0.00 | | |
| | Northern Indian Ocean | NA* | NA* | 0.00 | |
| | Indo-Australian Archipelago | 0.51 | 0.16 | NA* | 0.00 |
| Benthic Foraminifera | West Atlantic | 0.00 | | | |
| | Western Tethys | 0.21 | 0.00 | | |
| | Northern Indian Ocean | 0.48 | 0.30 | 0.00 | |
| | Indo-Australian Archipelago | 0.67 | 0.40 | 0.49 | 0.00 |

*Depositional environment difference between provinces*

The PBDB and literature based depositional-environment categories were consolidated into seven broader environmental groups based on their depositional setting. *Coastal environments* included coastal indeterminate, foreshore, shoreface, transition zone/lower shoreface, estuary/bay, and marginal-marine settings. *Deltaic environments* included delta front, delta plain, deltaic indeterminate, prodelta, interdistributary bay, and paralic settings. *Reef environments* included basin reefs, intrashelf/intraplatform reefs, perireef or subreef settings, platform/shelf-margin reefs, reef/buildup/bioherm settings, and slope/ramp reefs. *Shallow-marine environments* included carbonate indeterminate, lagoonal, lagoonal/restricted shallow subtidal, open shallow subtidal, peritidal, sand shoal, and shallow subtidal indeterminate settings. *Deep-marine* environments included deep-water indeterminate, slope, submarine-fan, and basinal carbonate, siliceous, and siliciclastic settings. *Offshore/shelf* environments included offshore, offshore indeterminate, offshore ramp, offshore shelf, deep subtidal ramp, deep subtidal shelf, and deep subtidal indeterminate settings. Occurrences classified as marine indeterminate, as well as those lacking depositional-environment information, were assigned to a marine-indeterminate category.

Depositional environments also varied among provinces and taxonomic groups across the Cenozoic (Fig. S6). We used a Bray-Curtis dissimilarity with a threshold of 0.5 (0 indicate greater similarity and values closer to 1 indicate dissimilarity) to identify province pairs with substantially different depositional environment composition (Table S2). Mollusca showed relatively similar depositional-environment composition among the provinces, with dissimilarity values of 0.25-0.28 where comparisons could be calculated. Similarly, Cnidaria showed low dissimilarity among the comparable province pairs, ranging from 0.13 to 0.16.

In contrast, benthic Foraminifera showed substantially greater differences in depositional-environment composition among provinces. Bray-Curtis dissimilarities for West Atlantic-Northern Indian Ocean (0.73), Western Tethys-Northern Indian Ocean (0.61), West Atlantic-Indo-Australian Archipelago (0.58), and Western Tethys-Indo-Australian Archipelago (0.63). The relatively high dissimilarities among Benthic Foraminifera indicate strong differences in the depositional environments could affect the diversity patterns in benthic foraminiferal hotspots.

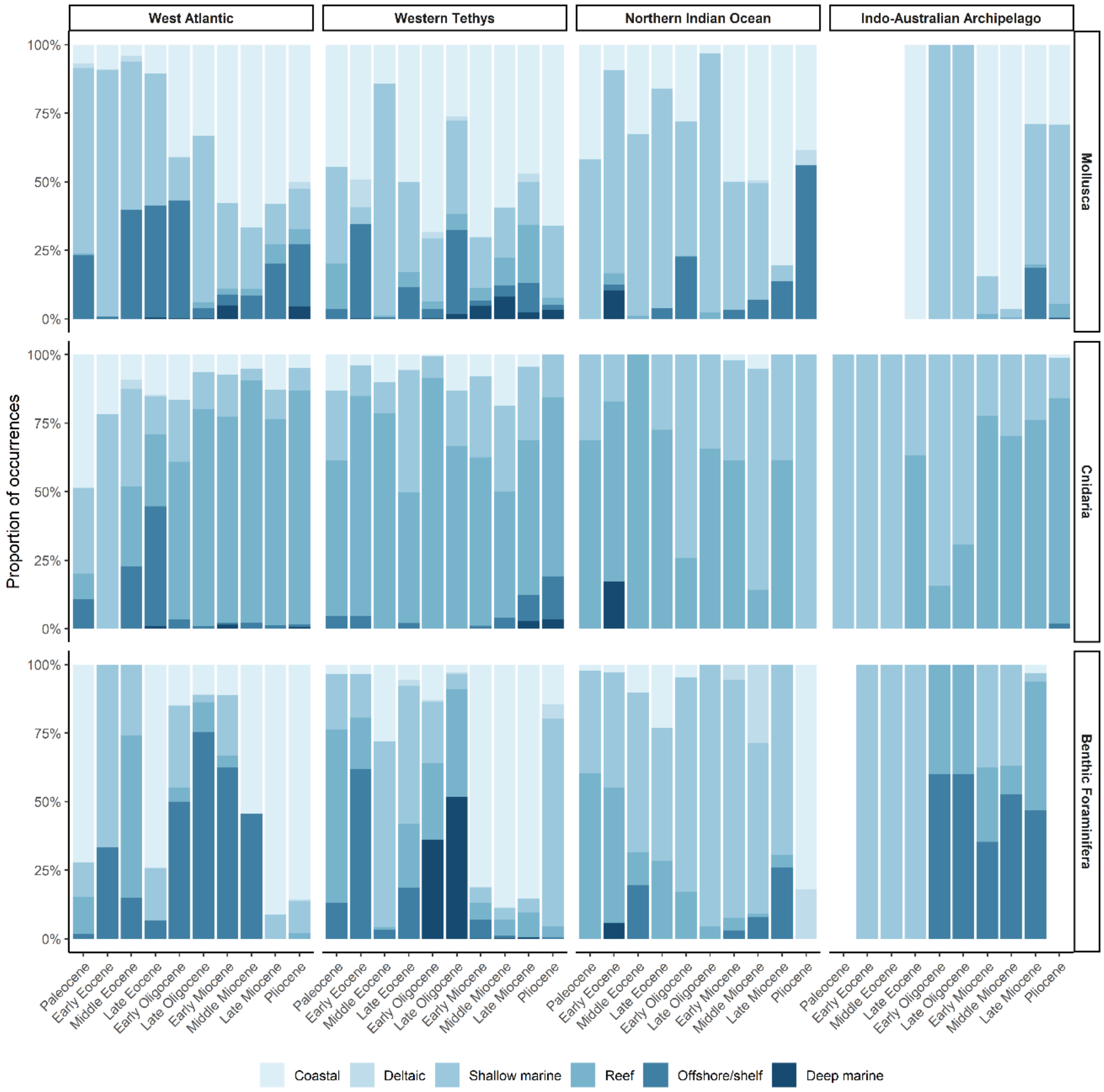


**Figure S6.** Depositional-environment composition of fossil occurrences across Cenozoic marine biodiversity provinces. Proportions of occurrences assigned to coastal, deltaic, shallow-marine, reef, offshore/shelf, and deep-marine environments are shown for Mollusca, Cnidaria, and benthic Foraminifera across the four biodiversity provinces and successive Cenozoic epochs. Marine-indeterminate occurrences are excluded from the proportional calculation.

**Table S2.** Bray-Curtis dissimilarity matrices showing differences in the composition of depositional environments among the four biogeographic provinces for Mollusca, Cnidaria, and Benthic Foraminifera across the Cenozoic. Marine-indeterminate occurrences were excluded from the analysis. *Cnidaria and Mollusca were removed from Northern Indian Ocean.

| **Taxon** | | West Atlantic | Western Tethys | Northern Indian Ocean | Indo-Australian Archipelago |
|---|---|---|---|---|---|
| Mollusca | West Atlantic | 0.00 | | | |
| | Western Tethys | 0.28 | 0.00 | | |
| | Northern Indian Ocean | NA* | NA* | 0.00 | |
| | Indo-Australian Archipelago | 0.25 | 0.28 | NA* | 0.00 |
| Cnidaria | West Atlantic | 0.00 | | | |
| | Western Tethys | 0.13 | 0.00 | | |
| | Northern Indian Ocean | NA* | NA* | 0.00 | |
| | Indo-Australian Archipelago | 0.14 | 0.16 | NA* | 0.00 |
| Benthic Foraminifera | West Atlantic | 0.00 | | | |
| | Western Tethys | 0.15 | 0.00 | | |
| | Northern Indian Ocean | 0.73 | 0.61 | 0.00 | |
| | Indo-Australian Archipelago | 0.58 | 0.63 | 0.43 | 0.00 |

### S2. Standardization of macroevolutionary processes

We quantified three components of diversity dynamics (origination, extinction, and migration) for each taxon, province, and time bin. These components were measured using genus-level First Appearance Datum (FAD) and Last Appearance Datum (LAD) records, with genus ranges estimated from Phanerozoic fossil occurrence data. Global FAD and LAD ranges were used to identify origination and extinction events for each time bin. We defined the components as follows:

(1) origination events: the number of genera whose FAD occurred within a given province and time bin;

(2) extinction events: the number of genera whose LAD occurred within a given province and time bin;

(3) migration events: the first occurrence of a genus in a province other than its inferred province of origin.

Raw counts of origination, extinction, and migration are confounded by uneven fossil sampling effort among biogeographic provinces. Therefore, to minimise this effect of sampling bias, we applied two independent resampling procedures and compared these results by correlating with raw occurrence counts.

#### *S2.1. Resampling the occurrence data using Coverage-quorum (q-value) resampling*

We resampled the fossil occurrence data using the same taxon-specific sample-coverage quorum (Good's $u$) previously determined for Shareholder Quorum Subsampling (SQS) of genus richness. For each iteration, occurrence records were resampled with replacement independently within each of the four focal provinces (West Atlantic, Western Tethys, Northern Indian Ocean, and Indo-Australian Archipelago) until the corresponding taxon-specific coverage quorum was reached. The four province-level resampled datasets were then combined, and each occurrence was classified according to the three macroevolutionary processes considered in this study: origination, extinction, and migration. This procedure was repeated for 1000 iterations. For each taxon, province, epoch, and macroevolutionary process, we calculated the mean and standard deviation of the process counts across the 1000 resampling iterations. The mean process count was then divided by the total number of raw fossil occurrences in the corresponding province and epoch to obtain an occurrence-standardized process rate.

To evaluate whether coverage-quorum resampling reduced the influence of variation in fossil sampling intensity, we tested the relationship between the standardized rates of macroevolutionary processes and the original number of fossil occurrences. Pearson correlation coefficients (*r*) were calculated separately for origination, extinction, and migration for each taxon-province combination, using the standardized process rate as the response variable and the total number of raw fossil occurrences per epoch as the predictor (Table S3).

A significant correlation between fossil occurrence abundance and a standardized process rate indicates that the estimated macroevolutionary process remains systematically associated with sampling intensity, suggesting that coverage-quorum resampling has not completely removed the influence of uneven fossil sampling for that taxon-province-process combination. In contrast, a non-significant correlation indicates that the standardized process rate does not show a detectable relationship with the number of raw fossil occurrences across epochs, suggesting that the coverage-quorum resampling has effectively reduced the detectable influence of sampling intensity on that process estimate.

*S2.2. Resampling the occurrence data using minimum occurrence rarefaction resampling*

In this method we identified, for each taxon, the province with the fewest fossil occurrences among the four focal provinces for entire Cenozoic, and used its occurrence count as a common sample size (n_min). In each iteration, we drew n_min number of rows with replacement, independently from each of the four focal provinces, ensuring all four provinces contributed an identical number of occurrences regardless of their original sampling intensity. The four province-level resampled datasets were then combined, and occurrences were classified according to the three macroevolutionary processes considered in this study: origination, extinction, and migration. This procedure was repeated for 1000 iterations. For each taxon, province, epoch, and macroevolutionary process, the mean and standard deviation of the process counts were calculated across the 1000 iterations. The mean process count was divided by the total number of raw fossil occurrences in the corresponding province and epoch to obtain an occurrence-standardized process rate.

To evaluate whether minimum-occurrence rarefaction reduced the influence of uneven fossil sampling, we tested the relationship between the standardized macroevolutionary process rates and the original number of fossil occurrences. Pearson correlation coefficients (*r*) were calculated separately for origination, extinction, and migration for each taxon-province combination, using the standardized process rate as the response variable and the total number of raw fossil occurrences per epoch as the predictor (Table S3).

A significant correlation indicates that the estimated macroevolutionary process rate remains systematically related to the original sampling intensity, suggesting that minimum-occurrence rarefaction has not completely removed the influence of uneven fossil sampling for that taxon-province-process combination. A non-significant correlation indicates that the standardized process rate has no detectable relationship with the original number of fossil occurrences across epochs, suggesting that minimum-occurrence rarefaction has reduced the detectable influence of sampling intensity on that process estimate.

Table S3: Pearson correlations between raw fossil occurrence counts and macroevolutionary processes calculated using the coverage-quorum (q-value) resampling and minimum-occurrence rarefaction resampling methods. Values are presented for three macroevolutionary processes- origination, extinction, and Immigration for each province. r denotes the Pearson correlation coefficient, and p denotes the corresponding significance level. Significant correlations ($p < 0.05$) are indicated in red.

| **Taxon** | **Province** | **Macro-evolutionary Process** | **Coverage-quorum** | | **Minimum-occurrence rarefaction** | |
|---|---|---|---|---|---|---|
| | | | **r** | **p value** | **r** | **p value** |
| Mollusca | West Atlantic | Origination | -0.68 | 0.03 | -0.57 | 0.08 |
| | | Extinction | 0.32 | 0.37 | 0.5 | 0.14 |
| | | Immigration | -0.65 | 0.04 | -0.68 | 0.03 |
| | Western Tethys | Origination | -0.48 | 0.16 | -0.44 | 0.2 |
| | | Extinction | -0.63 | 0.05 | -0.61 | 0.06 |
| | | Immigration | -0.44 | 0.2 | -0.58 | 0.08 |
| | Indo-Australian Archipelago | Origination | -0.95 | 0.05 | -0.93 | 0.07 |
| | | Extinction | 0.87 | 0.13 | 0.89 | 0.11 |
| | | Immigration | -0.66 | 0.34 | -0.59 | 0.41 |

| | | | | | | |
|---|---|---|---|---|---|---|
| Cnidaria | West Atlantic | Origination | -0.79 | 0.01 | -0.67 | 0.03 |
| | | Extinction | -0.28 | 0.43 | 0.05 | 0.89 |
| | | Immigration | -0.75 | 0.01 | -0.75 | 0.01 |
| | Western Tethys | Origination | -0.58 | 0.08 | -0.53 | 0.11 |
| | | Extinction | -0.69 | 0.03 | -0.63 | 0.05 |
| | | Immigration | -0.67 | 0.03 | -0.77 | 0.01 |
| | Indo-Australian Archipelago | Origination | -0.4 | 0.6 | -0.38 | 0.62 |
| | | Extinction | -0.76 | 0.25 | -0.76 | 0.24 |
| | | Immigration | -0.25 | 0.75 | -0.32 | 0.68 |
| Benthic Foraminifera | West Atlantic | Origination | 0.28 | 0.64 | 0.17 | 0.79 |
| | | Extinction | -0.29 | 0.63 | -0.37 | 0.54 |
| | | Immigration | -0.55 | 0.33 | -0.54 | 0.34 |
| | Western Tethys | Origination | -0.55 | 0.1 | -0.43 | 0.21 |
| | | Extinction | -0.7 | 0.02 | -0.63 | 0.05 |
| | | Immigration | -0.61 | 0.06 | -0.72 | 0.02 |
| | Northern Indian Ocean | Origination | 0.37 | 0.3 | 0.19 | 0.59 |
| | | Extinction | -0.14 | 0.7 | -0.17 | 0.64 |
| | | Immigration | -0.22 | 0.55 | -0.01 | 0.98 |
| | Indo-Australian Archipelago | Origination | NA | NA | NA | NA |
| | | Extinction | NA | NA | NA | NA |
| | | Immigration | NA | NA | NA | NA |
| Planktic Foraminifera | West Atlantic | Origination | -0.49 | 0.16 | -0.48 | 0.16 |
| | | Extinction | -0.55 | 0.1 | -0.55 | 0.1 |
| | | Immigration | -0.41 | 0.24 | -0.44 | 0.2 |
| | Western Tethys | Origination | -0.45 | 0.19 | -0.45 | 0.2 |
| | | Extinction | -0.55 | 0.1 | -0.5 | 0.14 |
| | | Immigration | -0.27 | 0.45 | -0.28 | 0.43 |
| | Northern Indian Ocean | Origination | -0.42 | 0.23 | -0.42 | 0.22 |
| | | Extinction | -0.53 | 0.12 | -0.54 | 0.11 |
| | | Immigration | -0.47 | 0.17 | -0.46 | 0.18 |
| | Indo-Australian Archipelago | Origination | 0.62 | 0.38 | 0.62 | 0.38 |
| | | Extinction | -0.21 | 0.79 | -0.22 | 0.78 |

| | | | | | | |
|---|---|---|---|---|---|---|
| | | Immigration | -0.86 | 0.14 | -0.85 | 0.15 |

Both standardization approaches reduced, but did not completely eliminate, correlations between raw occurrence counts and estimated macroevolutionary process rates. Under q-value resampling, significant negative correlations were detected in seven taxon-province-process combinations, primarily involving Mollusca and Cnidaria in the West Atlantic and Western Tethys, as well as extinction in benthic foraminifera in the Western Tethys (Table S3). Minimum-occurrence rarefaction produced five significant negative correlations, with a broadly similar concentration in the West Atlantic and Western Tethys (Table S3).

We proceeded with coverage-quorum (q-value) resampling as the preferred standardization approach because it accounts for differences in sampling completeness among provinces rather than equalizing provinces solely on the basis of their total number of fossil occurrences. Minimum-occurrence rarefaction standardizes the absolute number of sampled occurrences among provinces, but can substantially reduce the available information from more intensively sampled provinces and does not directly account for differences in taxonomic sampling completeness. Coverage-quorum resampling also provides consistency with the taxon-specific q values used in the Shareholder Quorum Subsampling (SQS) analysis, from which the standardized diversity curves were generated.

*S2.3. Dominant macroevolutionary processes operating in hotspots*

We evaluated the relative contributions of three macroevolutionary processes—origination, migration, and extinction—to the formation and maintenance of marine biodiversity hotspots. For each taxon, province, and epoch in which a province was identified as a hotspot, we compared the standardized rates of the three processes and identified the process with the highest rate as the dominant macroevolutionary process for that interval (Fig. S7). The resulting classification allowed us to determine whether hotspot intervals were predominantly associated with origination, migration, or extinction, and to compare the dominant processes through time and among the four biogeographic provinces.

Because the process rates were standardized for sampling intensity, comparisons were based on the relative contribution of each macroevolutionary process rather than differences in the absolute number of fossil occurrences or taxa among provinces.

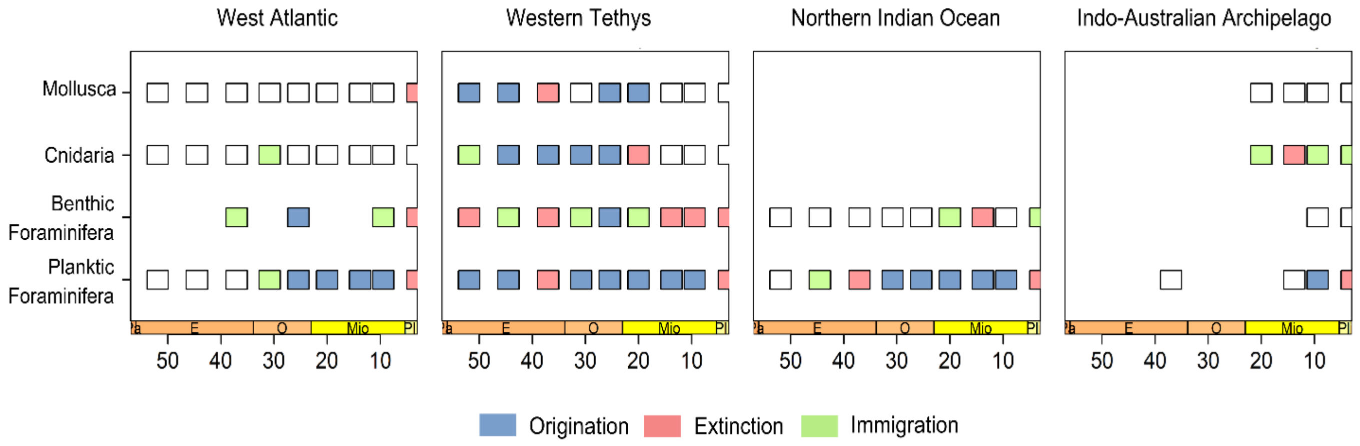


**Figure S7.** Classification of biodiversity hotspots based on dominant macroevolutionary processes across marine groups and provinces. Hotspots are classified for four marine groups-Mollusca, benthic foraminifera, planktic foraminifera, and Cnidaria—across Cenozoic time bins and biogeographic provinces. Each coloured box indicates the dominant process contributing to hotspot status within a given interval: origination (blue), immigration (green), or extinction (red).

S3. Evaluating environmental variation across hotspot and non-hotspot region

*S3.1. Sea surface temperature*

Cenozoic SST estimates were obtained from the compiled various paleo-proxies derived from the PhanSST database (Judd et al., 2022). Epoch-specific SST values were joined to the biodiversity dataset using the Cenozoic epoch designation. Because the provinces span different climatic zones, SST was assigned according to the climatic setting of each province. The tropical SST estimate was used for the West Atlantic, Northern Indian Ocean, and Indo-Australian Archipelago, whereas the temperate SST estimate was used for the Western Tethys. Global SST estimates were not used in the subsequent analyses. Consequently, SST represents the estimated thermal conditions characteristic of the climatic zone occupied by each province during each epoch.

*S3.2. Change in regional shelf area*

Palaeoshelf area was estimated from the 1° × 1° palaeogeographic digital elevation models (PaleoDEMs) of Scotese and Wright (2018). NetCDF palaeogeographic grids representing 0, 10, 15, 20, 25, 30, 35, 45, 50, and 60 Ma were used to represent the Pliocene, Late Miocene, Middle Miocene, Early Miocene, Late Oligocene, Early Oligocene, Late Eocene, Middle Eocene, Early Eocene, and Paleocene, respectively. For each biodiversity province, the corresponding latitude-longitude bounding box was extracted from each palaeogeographic grid. Grid cells with palaeoelevation values between −1000 and 0 m were classified as shelf environments. Palaeoshelf area was calculated as the number of qualifying 1° × 1° grid cells multiplied by an approximate cell area of 111 × 111 $km^2$ (valid for tropical regions). The resulting values represent the total estimated shelf area within each predefined province for each time interval. This approach provides a consistent measure of the spatial availability of shallow-marine habitat through time.

*S3.3. Global mean sea level*

Global sea-level estimates were obtained from the smoothed sea-level curve of Miller et al. (2020). Individual sea-level observations were assigned to Cenozoic epochs using standard epoch boundaries. To obtain a representative sea-level value for each epoch, we defined a temporal midpoint for each epoch and calculated the mean smoothed sea-level value of all observations occurring within ±1 Myr of that midpoint. The resulting mean value was used as the epoch-specific sea-level covariate and was assigned to all provinces within that epoch.

Thus, sea level represents a global, time-varying environmental variable, rather than a province-specific measurement.

### S4. Causal pathway analysis of environmental and macroevolutionary drivers of Cenozoic Hotspot Dynamics

We quantified the causal pathways underlying hotspot formation by integrating environmental drivers and macroevolutionary processes using Bayesian path analysis implemented in the *brms* package in R (Bürkner, 2017). The model specified directional relationships among environmental variables (sea surface temperature, mean sea level, and shelf area) to macroevolutionary processes (origination, extinction, and migration), and finally to hotspot occurrence across marine clades.

#### *S4.1. Direct effects*

Direct effects represent the immediate influence of macroevolutionary processes on hotspot occurrence and were estimated as posterior effect sizes of origination, extinction, and migration on hotspot probability while accounting for other variables in the model (Table S4).

**Table S4**. Direct effects of macroevolutionary processes on hotspot dynamics estimated using Bayesian path analysis (brms). Mean posterior estimates and 95% credible intervals are shown for origination, extinction, and migration across taxa. Significant effects are indicated where credible intervals do not overlap zero.

| **Taxon** | **Process** | **Mean posterior estimates** | **Q2.5** | **Q97.5** | **Significant** |
|---|---|---|---|---|---|
| Mollusca | Origination | 1.56 | 0.27 | 3.34 | TRUE |
| | Extinction | 1.14 | -0.34 | 3.25 | FALSE |
| | Immigration | -0.87 | -2.80 | 0.42 | FALSE |
| Cnidaria | Origination | 2.23 | 0.34 | 4.92 | TRUE |
| | Extinction | 0.58 | -0.84 | 2.13 | FALSE |
| | Immigration | -0.44 | -1.70 | 0.93 | FALSE |
| Benthic Foraminifera | Origination | 0.43 | -3.37 | 3.83 | FALSE |
| | Extinction | 0.39 | -2.17 | 2.43 | FALSE |
| | Immigration | -1.08 | -3.32 | 0.58 | FALSE |

| | | | | | |
|---|---|---|---|---|---|
| Planktic Foraminifera | Origination | 0.43 | -3.37 | 3.83 | FALSE |
| | Extinction | 0.39 | -2.17 | 2.43 | FALSE |
| | Immigration | -1.08 | -3.32 | 0.58 | FALSE |

*S4.2. Indirect effects*

Indirect effects capture mediated pathways through which environmental variables influence hotspot occurrence via their effects on macroevolutionary processes. These effects were quantified by multiplying posterior estimates along each causal pathway (Table S5).

**Table S5.** Effects of environmental variables on macroevolutionary processes across marine taxa estimated using Bayesian path analysis (brms). Mean posterior estimates and 95% credible intervals are shown for the effects of Sea Surface Temperature (SST), shelf area, and mean sea level on origination, extinction, and migration. Significant effects are indicated where credible intervals do not overlap zero.

| Taxon | Environment | Process | Mean posterior estimates | Q2.5 | Q97.5 | Significant |
|---|---|---|---|---|---|---|
| Mollusca | SST | Origination | 0.10 | -0.28 | 0.66 | FALSE |
| | | Extinction | -0.21 | -0.84 | 0.10 | FALSE |
| | | Immigration | 0.03 | -0.33 | 0.44 | FALSE |
| | Shelf Area | Origination | 0.93 | 0.13 | 2.28 | TRUE |
| | | Extinction | 0.25 | -0.15 | 1.08 | FALSE |
| | | Immigration | 0.16 | -0.21 | 0.80 | FALSE |
| | Mean sea level | Origination | 0.44 | 0.01 | 1.26 | TRUE |
| | | Extinction | 0.04 | -0.44 | 0.55 | FALSE |
| | | Immigration | -0.05 | -0.59 | 0.43 | FALSE |
| Cnidaria | SST | Origination | 0.01 | -0.62 | 0.62 | FALSE |
| | | Extinction | -0.12 | -0.60 | 0.18 | FALSE |
| | | Immigration | 0.04 | -0.19 | 0.36 | FALSE |
| | Shelf Area | Origination | 1.03 | 0.11 | 2.67 | TRUE |
| | | Extinction | 0.13 | -0.22 | 0.69 | FALSE |
| | | Immigration | 0.08 | -0.25 | 0.51 | FALSE |

| | | | | | | |
|---|---|---|---|---|---|---|
| | Mean sea level | Origination | 0.63 | 0.02 | 1.84 | TRUE |
| | | Extinction | 0.06 | -0.21 | 0.44 | FALSE |
| | | Immigration | -0.13 | -0.69 | 0.30 | FALSE |
| Benthic Foraminifera | SST | Origination | 0.01 | -0.51 | 0.55 | FALSE |
| | | Extinction | -0.06 | -0.60 | 0.48 | FALSE |
| | | Immigration | 0.09 | -0.36 | 0.69 | FALSE |
| | Shelf Area | Origination | 0.13 | -1.51 | 1.84 | FALSE |
| | | Extinction | 0.01 | -0.73 | 0.75 | FALSE |
| | | Immigration | 0.26 | -0.38 | 1.23 | FALSE |
| | Mean sea level | Origination | 0.10 | -0.94 | 1.15 | FALSE |
| | | Extinction | 0.06 | -0.58 | 0.74 | FALSE |
| | | Immigration | -0.21 | -1.12 | 0.47 | FALSE |
| Planktic Foraminifera | SST | Origination | 0.01 | -0.51 | 0.55 | FALSE |
| | | Extinction | -0.06 | -0.60 | 0.48 | FALSE |
| | | Immigration | 0.09 | -0.36 | 0.69 | FALSE |
| | Shelf Area | Origination | 0.13 | -1.51 | 1.84 | FALSE |
| | | Extinction | 0.01 | -0.73 | 0.75 | FALSE |
| | | Immigration | 0.26 | -0.38 | 1.23 | FALSE |
| | Mean sea level | Origination | 0.10 | -0.94 | 1.15 | FALSE |
| | | Extinction | 0.06 | -0.58 | 0.74 | FALSE |
| | | Immigration | -0.21 | -1.12 | 0.47 | FALSE |
